# Biofilm-derived polymer networks constrain immune cell motility through adhesion-mediated physical interactions

**DOI:** 10.1101/2025.03.11.642532

**Authors:** Kaman Ma, Yeping Ma, Song Lin Chua

## Abstract

Understanding how immune cells navigate complex microbial environments requires models that integrate biophysical and biological realism. However, the microscale mechanisms underlying phagocyte migration on pathogenic biofilms remain poorly understood, and there remains a lack of experimental models for study of 3D-interactions between host cells and biofilms. Here, we present a multiscale microfluidic secondary-infection-on-a-chip incorporating human-cell-based model or an ex vivo perfused porcine skin system, to study macrophage migration across bacterial biofilms under controlled flow and spatial confinement. Using *Pseudomonas aeruginosa* as model pathogen, we demonstrated that biofilm matrix components, specifically the exopolysaccharide Psl, physically impede macrophage motility and directional persistence. Exopolysaccharide-deficient mutants allowed near-normal macrophage migration. Persistent random walk (PRW) simulation modeling recapitulated these motility dynamics, revealing altered migration coefficients within biofilm matrices. Consistent with cell-based model, biofilms impede macrophages on the skin tissue. Hence, our results establish an experimental and modelling framework for studying physical constraints in host-pathogen interactions, providing new insights into mechanical and spatial barriers that shape immune responses to biofilm infections.

## Introduction

Bacteria form multicellular biofilms encased in self-produced exopolymeric substances, as a form of protection from their predators in diverse habitats, such as natural environment and human infections (1). The predators range from phagocytic eukaryotes (immune cells and amoeba) and bacterivorous animals, to bacteriophages and predatory bacteria. Protected by a complex matrix of extracellular polymeric substances (EPS) comprising of varying compositions of exopolysaccharides, adhesion proteins and extracellular DNA (eDNA) (2), resident microbes can prevent predation by their predators (3, 4).

The direct interaction between phagocytic cells and biofilms plays a key role in chronic infections that pose a significant challenge to human health. During the initial stage of infection, frontline resident macrophages act as first responders from the innate immunity (5), which migrate towards the site of infection to combat microbial invasion (6). While interactions between phagocytes and single-cell planktonic bacteria are well established, there are few studies on phagocytic eukaryotes-biofilm interactions. From the biochemical point-of-view, the chemical interactions between the macrophages and *Pseudomonas aeruginosa* biofilms during infections have been previous described (7, 8). Physical studies had shown that macrophages and amoebas often encounter ‘frustrated phagocytosis’, as they cannot phagocytose larger and stickier bacterial aggregates (9, 10). However, it is unclear if phagocytic cell motility and migration can be altered by presence of biofilms. Examination of both model organisms’ physical interactions could reveal novel mechanistic insights on phagocyte-biofilm interactions.

The lack of suitable coculture models also contributed to the poor understanding of phagocytic cell-biofilm interactions (11), as they do not simulate the migration to and motility of the phagocytic cell on the biofilms. Moreover, most studies employ 2D cellular monolayers which cannot recreate the 3D positioning of cells in a physiological environment. To address the shortcomings of 2D cultures, the use of microfluidics enables the rapid development of biofilms within 12 hrs and introduction of eukaryotic cells into 3D-biofilm models to study host-biofilm interactions (12).

In parallel, the *ex vivo* porcine skin model provided a physiologically relevant environment, preserving native dermal collagen architecture, extracellular matrix density, and interstitial fluid dynamics that influence cell migration (13, 14). Porcine skin is particularly suitable for infection studies because of its structural and immunological similarity to human skin (15), including comparable epidermal thickness, dermal composition, and immune cell distribution (14). Moreover, *in vivo* animal models are cost- and resource-intensive, which lead to the shift of research using *in vivo* models to *ex vivo* models. Hence, this raises the need for experimental models which can bridge reductionist cells models and tissue-level complex ones, and model immune cell-biofilm interactions.

Here, we first develop a microfluidics-based 3D-phagocytic cell-biofilm interactions model under continuous flow, which simulates phagocytic cell migration to the biofilms and their movements on the biofilms. Using human macrophages as proof-of-concept, we report that *P. aeruginosa* biofilms could impede phagocytic cell movements. By screening the *P. aeruginosa* mutant library of known biofilm matrix components, we showed that Psl, a *P. aeruginosa* biofilm exopolysaccharide (16), could alter macrophage tracks and reduce their motility. Macrophages displayed ‘frustrated motility’ even as they expended ATPs and synthesized actin filaments after being trapped on the biofilms. Exogenous addition of Psl to biofilm-deficient *P. aeruginosa* mutant could restore the frustrated motility phenotype in macrophages.

We observed a striking concordance in macrophage behaviour in our *ex vivo* porcine model, where wild-type *P. aeruginosa* biofilms reduced macrophage motility, while the pro-biofilm mutant Δ*wspF* (17) almost completely arrested macrophage movement. These consistent results across two models strengthen the conclusion that Psl-rich biofilms can not only resist phagocytosis but also physically hinder immune cell migration within infected tissue. Hence, insights gained from this study not only advance our understanding of host-biofilm interactions but also offer promising avenues for the development of novel therapeutic interventions targeting biofilm-associated infections.

## Results

### Biofilms impede locomotion and restrict motility of phagocytic cells

Based on our previously described model (12), we first developed a microfluidic phagocytic cell-biofilm interactions model, where biofilms were cultivated in channels containing microwells followed by introduction of LPS-activated human macrophages into the flow chamber (**Figure 1a-b)**. This simulated a migratory behaviour of activated macrophages to the biofilms, leading to their attachment and motility on the biofilms. In the absence of biofilms, macrophages could roam the surface freely (**Figure 1c-d, Supplementary Video 1**), with velocities similar to previous work (18) (**Figure 1e**). However, *P. aeruginosa* PAO1 biofilms could restrict the motility and the distance travelled by macrophages (**Figure 1c-d, Supplementary Video 2**), where the macrophage velocities decreased (**Figure 1e**). Macrophages were completely immobilized when encountering a pro-biofilm mutant Δ*wspF* (**Figure 1c-e, Supplementary Video 3**). Mutation in the *wspF* gene will lead to de-repression of the *wspR* c-di-GMP synthesis protein and enhanced biofilm formation, which is a common trait in clinical isolates (19). This showed that biofilms could restrict the motility of macrophages, resulting in limited distance covered by the macrophages.

**Figure 1.**
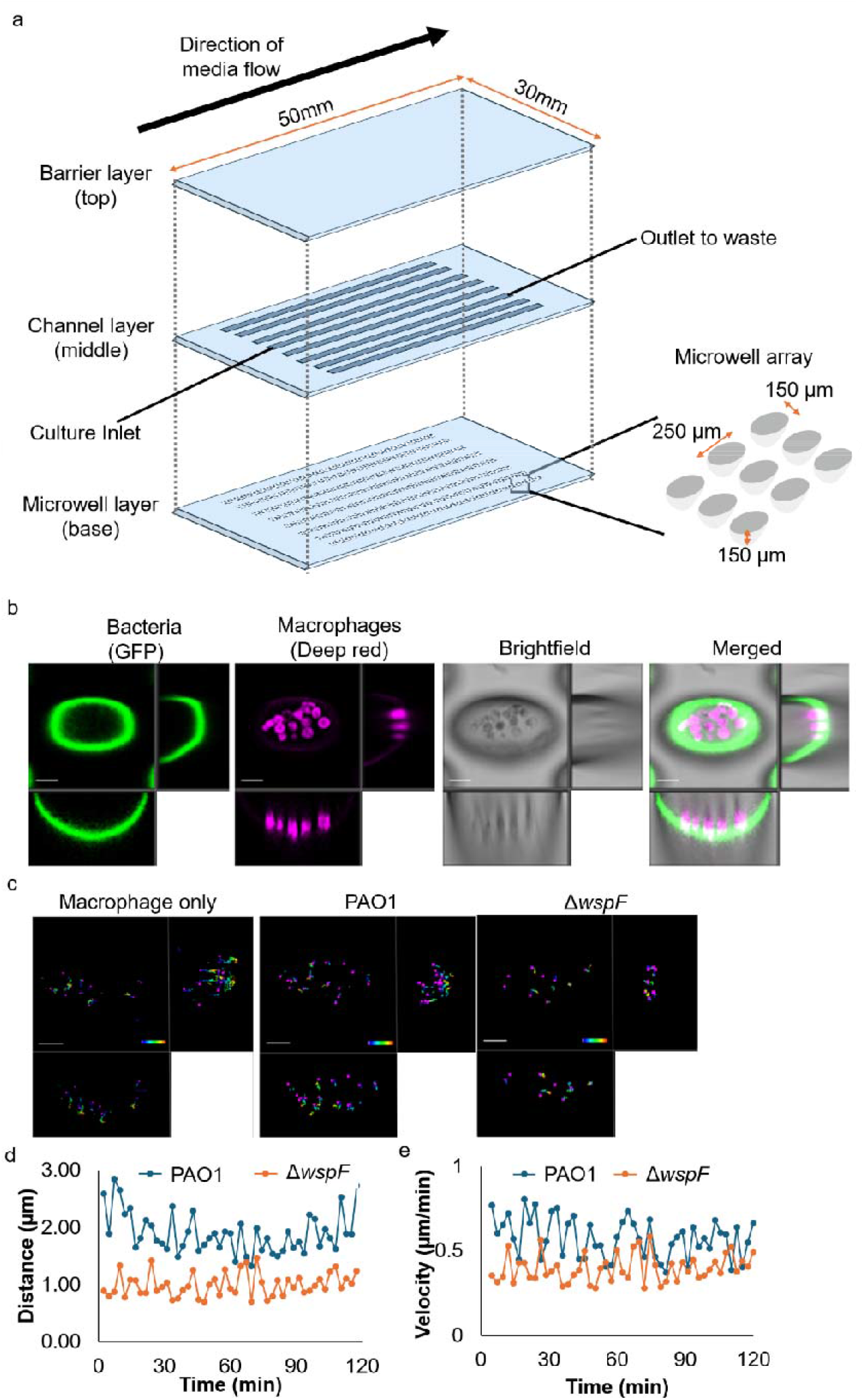
Biofilms impede macrophage motility. (a) Schematic diagram of microfluidics macrophage-biofilm model. (b) Representative CLSM images of macrophage-biofilm interactions model. Scale bar: 10 µm. (c) Representative tracks, (d) average distance and (e) average velocities of macrophages on PAO1 and Δ*wspF* mutant biofilms. Means and SD from triplicate experiments in three independent trials are shown. **P < 0.01; ***P < 0.001; n.s: not significant; One-Way ANOVA.

To show that the restriction of macrophages was not attributed to death or inactivity, we first quantified the death rate of the macrophages using propidium iodide (PI) and found that the macrophages remain mostly alive during their interactions with the biofilm (**Supplementary Figure 1a**). Moreover, macrophages remained consistently active, as they continued to expend energy in the form of ATP (**Supplementary Figure 1b**) and establish new actin filaments (**Supplementary Figure 1c**), both of which are crucial to macrophage motility (20, 21).

### Psl is more important than Pel at impeding phagocytic cell motility

Since *wspF* and its associated operon control the production of both Pel and Psl exopolysaccharides in biofilms, we seek to identify the biofilm matrix component which can restrict macrophage motility. To maximize the phenotypic effects of the specific exopolysaccharide and ensure our further observations were dependent solely on the specific exopolysaccharide, we introduced mutations in the EPS genes within the Δ*wspF* mutant. In this case, the presence of *wspF* mutation would boost the production of the exopolysaccharide whose synthesis genes were not mutated. Hence, unless specified otherwise, the Δ*wspF* would therein be referred to Pel^+^Psl^+^; Δ*wspF*Δ*pelA* was Pel^−^Psl^+^; Δ*wspF*Δ*pslBCD* was Pel^+^Psl^−^; Δ*wspF*Δ*pelA*Δ*pslBCD* was Pel^−^Psl^−^.

We showed that macrophages could roam more freely in the Pel^−^Psl^−^ biofilm which lost its Pel and Psl exopolysaccharides (**Figure 2a, Supplementary Video 4**), allowing them to move at higher velocity and longer distances (**Figure 2b-c**). Interestingly, macrophages remained constrained in Pel^−^Psl^+^ biofilm (**Figure 2a-c, Supplementary Video 5**), whereas Pel^+^Psl^−^ biofilm could not restrict macrophage motility (**Figure 2a-c, Supplementary Video 6**), indicating that the loss of Psl was deleterious to the biofilm at impeding macrophage motility. We also found qualitatively similar results for strains with non-*wspF*-mutated backgrounds (PAO1, Δ*pelA*, Δ*pslBCD*, and Δ*pelA*Δ*pslBCD* mutants) (**Supplementary Figure 2a-c, Supplementary Videos 2, 7-9**).

**Figure 2.**
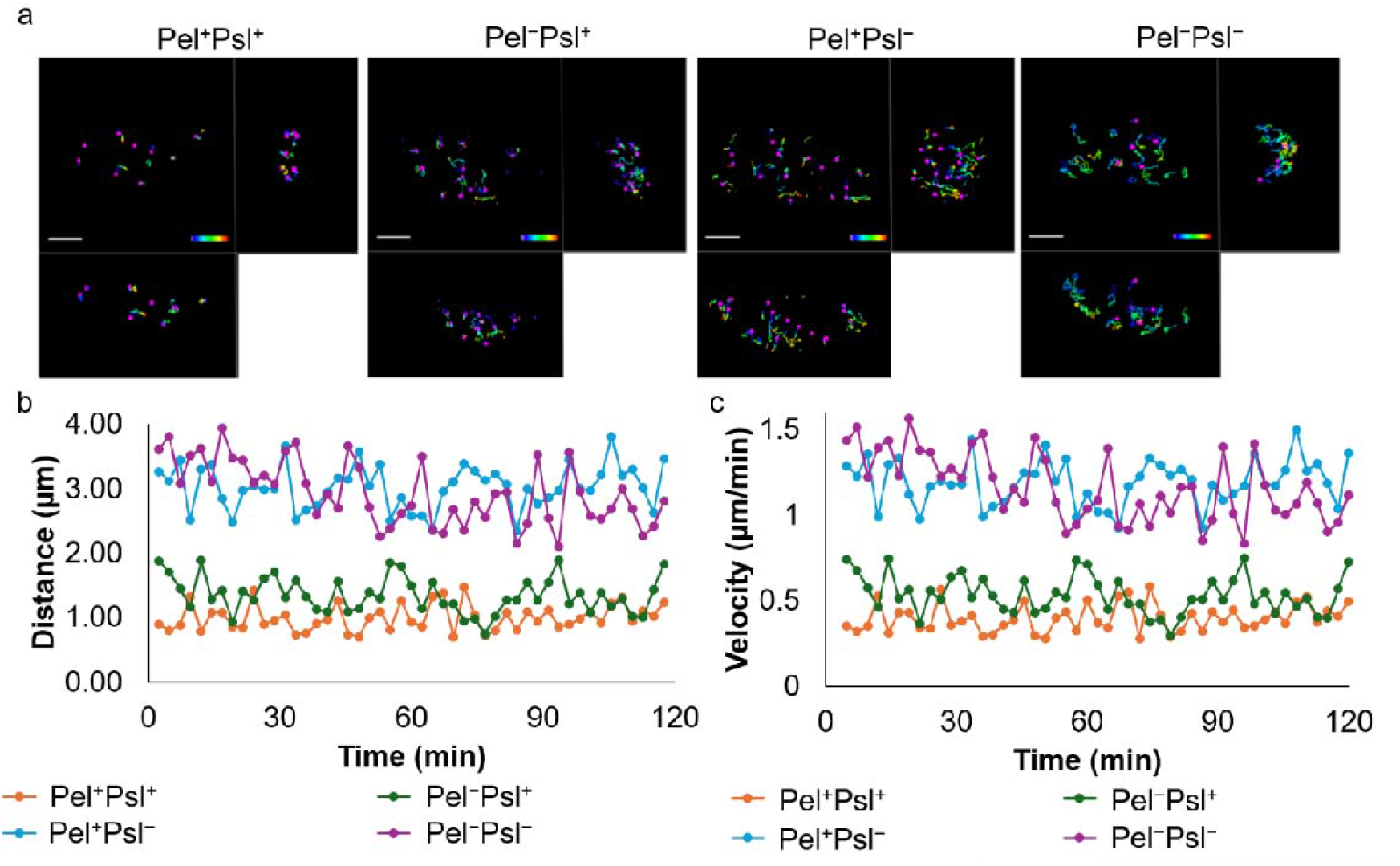
Psl is more important than Pel at impeding macrophage motility. (a) Representative tracks, (b) average distance, and (c) average velocity of macrophages on Δ*wspF* (Pel^+^Psl^+^), Δ*wspF*Δ*pelA* (Pel^−^Psl^+^), Δ*wspF*Δ*pslBCD* (Pel^+^Psl^−^), and Δ*wspF*Δ*pelA*Δ*pslBCD* (Pel^−^Psl^−^). Means and SD from triplicate experiments in three independent trials are shown. **P < 0.01; ***P < 0.001; n.s: not significant; One-Way ANOVA.

### Exogenous addition of Psl to biofilm-deficient mutant could impede phagocytic cell motility

To confirm that the physical presence of Psl was crucial to impeding macrophage motility, we coated 1 µg ml^-1^ of purified exogenous Pel or Psl polysaccharides on the surface, followed by cultivation of the Pel^−^Psl^−^ biofilm for observation of macrophage motility. While Pel-treated Pel^−^Psl^−^ could not effectively impede macrophage motility, macrophage motility was restricted by Psl-treated Pel^−^Psl^−^ biofilms (**Figure 3a-c, Supplementary Video 10-11**). As this concentration corroborated with the physiological levels of exopolysaccharides in the biofilms (22), this confirmed that physiologically-relevant Psl concentrations could enhance biofilm matrix and impede macrophage motility.

**Figure 3.**
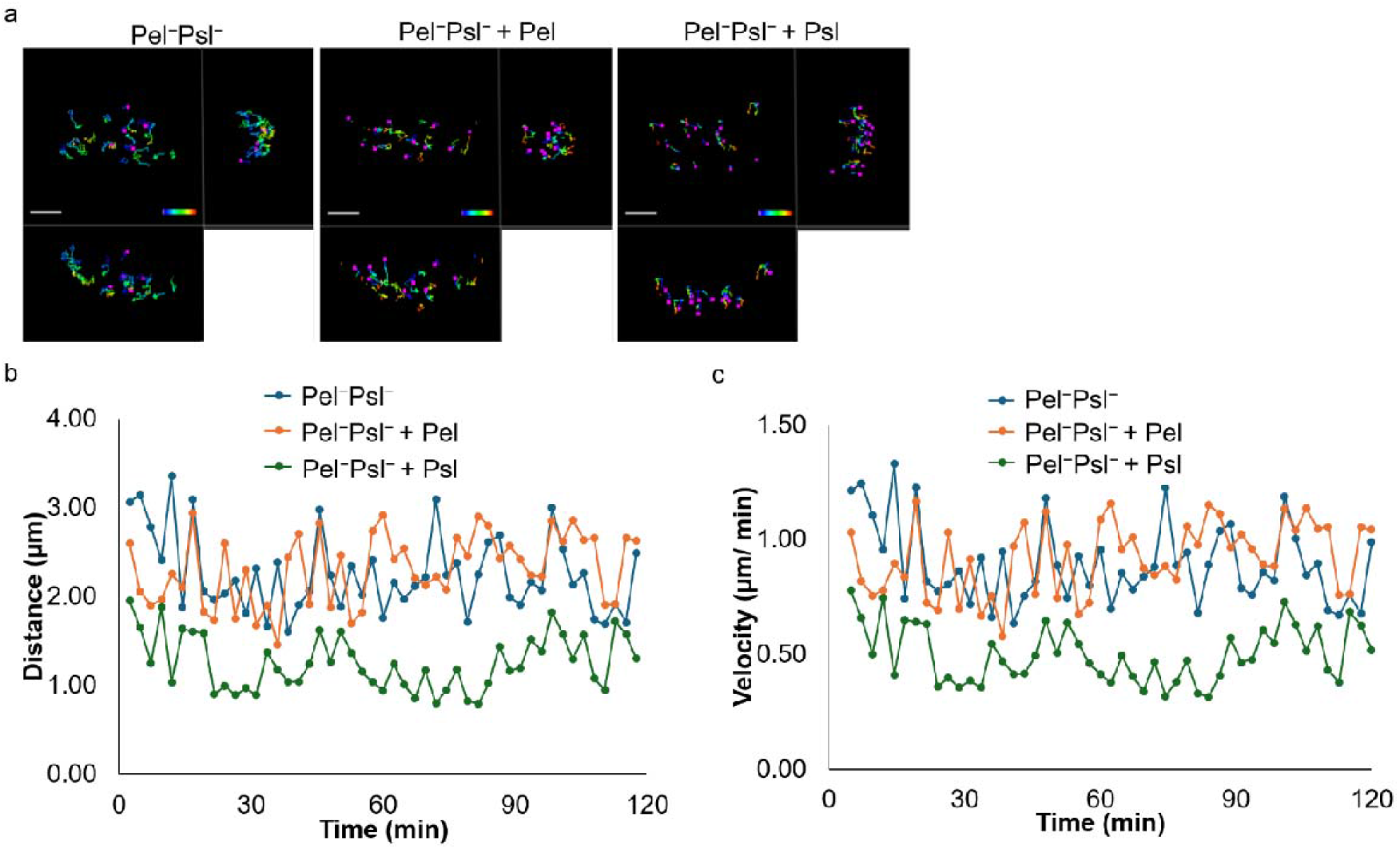
Exogenous addition of Psl to biofilm-deficient mutant could restore frustrated motility phenotype in macrophages. (a) Representative tracks, (b) average distance, and (c) average velocity of macrophages on Δ*wspF*Δ*pelA*Δ*pslBCD* (Pel^−^Psl^−^) on surfaces coated with Pel or Psl. Means and SD from triplicate experiments in three independent trials are shown. **P < 0.01; ***P < 0.001; n.s: not significant; One-Way ANOVA.

### Mathematical simulation validates phagocytic cell motility on biofilms in a persistent random walk (PRW) model

The PRW model is previously used to describe cell motility in 3D dimensions (23), especially for motile macrophages (24). Based on the macrophage tracks on different biofilms, our mathematical simulation showed that macrophages exhibited distinct biased persistent random walk patterns on different mutant biofilms (**Figure 4a-b**). Our heatmap further revealed that macrophages had similar behaviour on Pel^+^Psl^+^ and Pel^−^Psl^+^ across the entire duration of observation, which was distinct from on Pel^+^Psl^−^ and Pel^−^Psl^−^ (**Figure 4c**). This corroborated with our findings that Psl-containing biofilms could alter macrophage motility.

**Figure 4.**
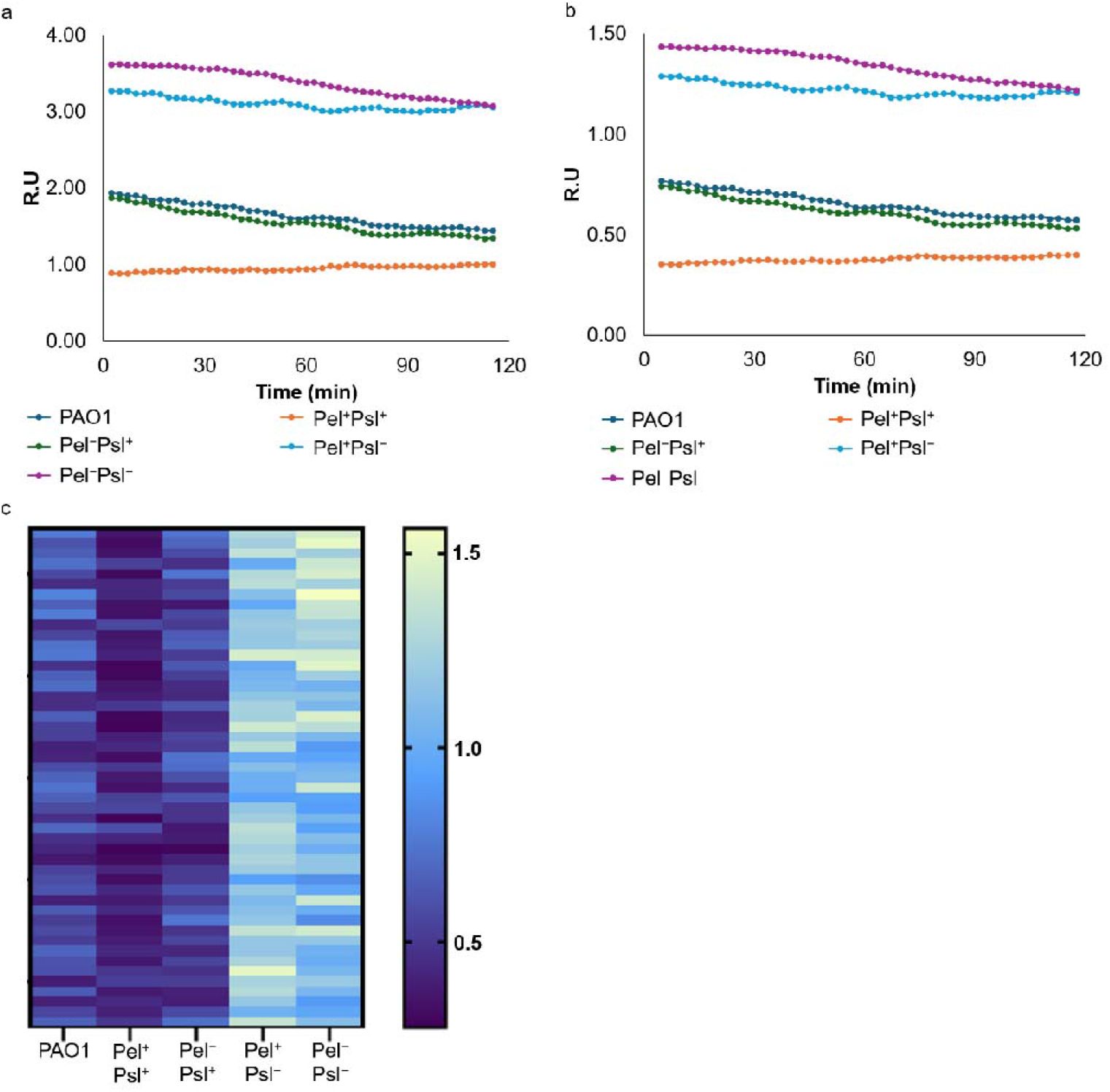
A PRW model of macrophage motility on biofilms. Mathematical simulation of (a) relative distance and (b) relative velocity of macrophages motility on various biofilm substrates. (c) Heatmap displaying differences in macrophages motility on various biofilm substrates.

### Ex vivo porcine biofilm infection model revealed poor macrophage motility on biofilms

To validate our microfluidics observations in a physiologically relevant setting, we established an *ex vivo* porcine skin biofilm infection model in which freshly excised dermal tissue was inoculated with *P. aeruginosa* strains to allow biofilm formation on the epidermal surface (**Figure 5a-b**). We could observe the macrophages on the wild-type PAO1 biofilms which was pre-established on the porcine skin (**Figure 5c**). In corroboration with our microfluidics results, wild-type PAO1 biofilms significantly reduced macrophage displacement and track length on the wounded skin, compared to Δ*pelA*Δ*pslBCD-*infected skin surfaces (**Supplementary Figure 3, Supplementary Video 12-13**). Our quantitative motility analysis showed a significant reduction in mean velocity and persistence in the presence of wild-type biofilms (**Supplementary Figure 3**).

**Figure 5.**
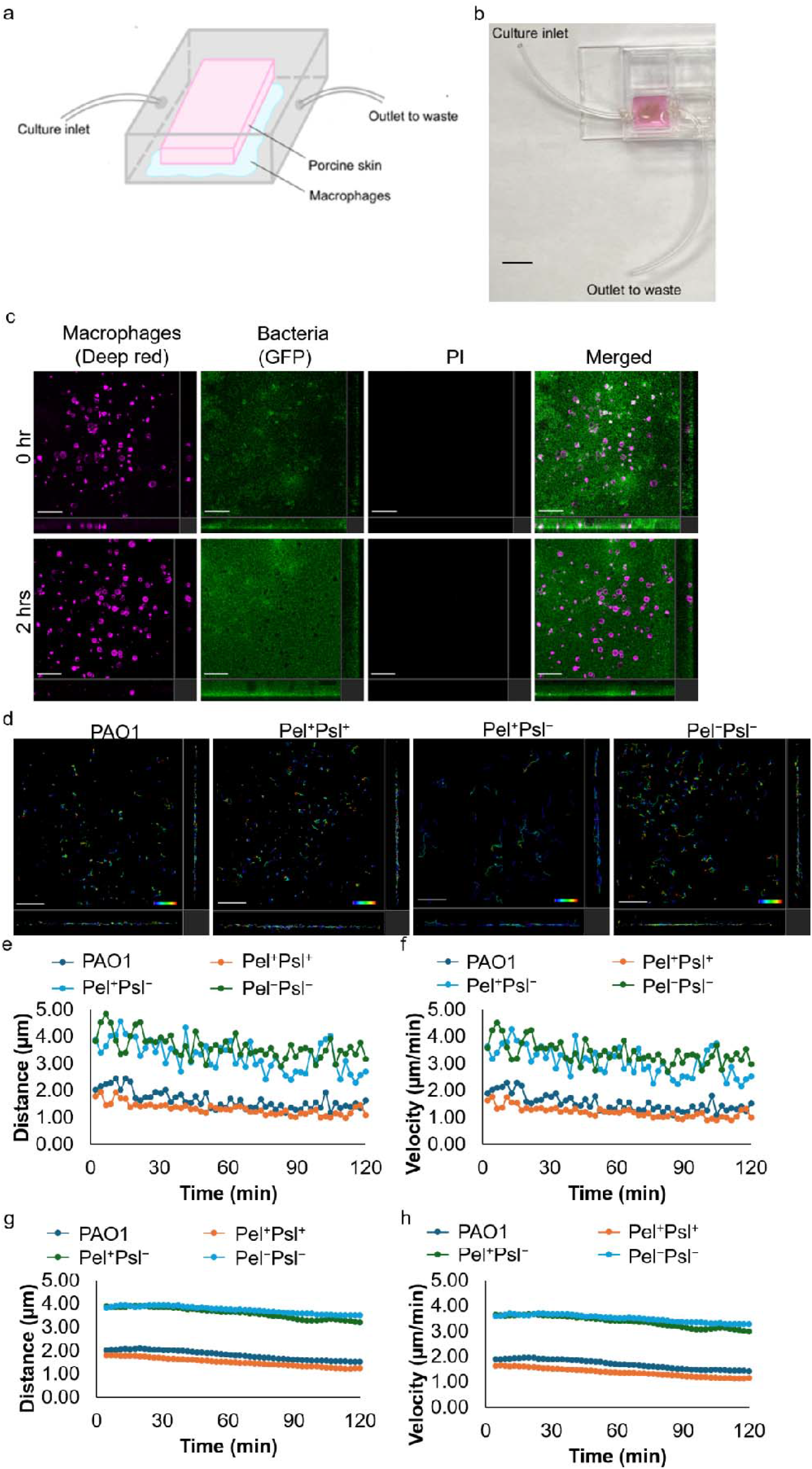
Macrophages exhibited low motility on *P. aeruginosa* biofilms in an *ex vivo* porcine skin infection model. (a) Schematic diagram and (b) representative image of the porcine skin biofilm infection model. Scale bar: 10 mm. (c) Representative CLSM images of macrophages on the wild-type PAO1 biofilms which was pre-established on the porcine skin. Scale bar: 10 µm. (d) Representative tracks, (e) average distance, and (f) average velocity of macrophages on PAO1, Δ*wspF* (Pel^+^Psl^+^), Δ*wspF*Δ*pslBCD* (Pel^+^Psl^−^), and Δ*wspF*Δ*pelA*Δ*pslBCD* (Pel^−^Psl^−^) biofilms pre-established on the porcine skin. Mathematical simulation of (g) relative distance and (h) relative velocity of macrophages motility on various biofilm pre-established on the porcine skin, based on PRW model. Means and SD from triplicate experiments in three independent trials are shown. **P < 0.01; ***P < 0.001; n.s: not significant; One-Way ANOVA.

Moreover, the Pel^+^Psl^+^ mutant with overproduction of EPS, almost completely abolished macrophage migration, with cells exhibiting minimal displacement from their point of entry (**Figure 5d-f, Supplementary Video 14-16**). In contrast, macrophages on tissue infected with the biofilm-deficient mutant lacking EPS (Pel^−^Psl^−^) had higher levels of migration than the Pel^+^Psl^+^-infected wound. Our PRW model also supported these results (**Figure 5g-h**), confirming that our observations are applicable in both biofilm-only and mammalian tissue-based models.

In summary, our findings confirmed that biofilm matrix components, particularly Psl, can physically hinder macrophage migration in a native tissue environment, mirroring the inhibitory effects observed in the engineered 3D microfluidics platform.

## Discussion

The biofilm matrix is an adhesive substrate with multiple functions, such as offering protection and metabolic exchanges of resident microbes (25). Other than preventing the effective phagocytosis of biofilm bacteria and enhancing immune evasion, our study using microfluidics reveals a novel function of the biofilm matrix restricting phagocytic cell motility, which highlights the profound property of biofilm adhesiveness on phagocytic cell behaviour. Exopolysaccharides, especially Psl, were responsible for impeding macrophage movement. By creating a highly adhesive environment, Psl likely enhances the strength of focal adhesions between the phagocytic cell and the biofilm matrix, making it difficult for the cells to move during migration. By constraining phagocytic cell within the confines of biofilm, this may facilitate bacterial evasion of predators in the environment, and immune cells in human infections.

Next, our study utilized the PRW model to analyze macrophage tracks on different *P. aeruginosa* biofilms, thereby revealing important insights into how biofilm matrix components influence immune cell behavior. The presence of Psl in biofilm matrix may create an adhesive environment that restricts macrophage movement, as reflected in the biased random walk patterns. These distinct motility patterns observed in our simulations are consistent with previous studies suggesting that macrophage motility can be influenced by the physical and chemical properties of their environment. Additionally, extending the PRW model to account for complex biofilm heterogeneity and macrophage signaling pathways would provide a more comprehensive understanding of immune cell-biofilm interactions.

Our *ex vivo* porcine skin infection model provided a crucial validation of these microfluidics-based observations under tissue-like conditions. Unlike reductionist *in vitro* systems, porcine skin preserves the extracellular matrix architecture, epidermal–dermal interface, and interstitial fluid dynamics that govern immune cell behaviour (26). Furthermore, using the *ex vivo* porcine skin model offers a valuable intermediate platform for preclinical testing of such anti-biofilm strategies (27), thus bridging the gap between *in vitro* mechanistic studies and *in vivo* infection models. We showed that macrophages encountering biofilm-covered tissue similarly exhibited severely impaired motility, particularly in the presence of Psl-overproducing Δ*wspF* biofilms. Moreover, we observed that the macrophages often adopted a rounded, sessile morphology rather than the elongated, polarized shape typical of actively migrating cells, implying that matrix components such as Psl may modify the surrounding tissue microenvironment and lead to altered cytoskeletal organization in macrophages.

The convergence of results from both systems (microfluidics system and porcine skin model) strengthens our conclusion that biofilm matrix components, especially Psl, represent a potent and generalizable barrier to immune cell access. Clinically, our findings have significant implications for chronic wound infections, where *P. aeruginosa* biofilms are frequently encountered and immune clearance is often incomplete. By impeding macrophage migration, biofilms may prolong bacterial persistence and exacerbate tissue damage. Targeting biofilm matrix integrity, particularly Psl, with an antibiofilm agent (12, 28) may therefore enhance immune cell penetration and pathogen clearance.

From an ecological perspective, our findings can be viewed as an extension of predator–prey interactions, where the biofilm serves as a barrier that shields bacteria from engulfment but also as a terrain modifier that limits predator mobility. Like macrophages, amoebae rely on active motility to locate and engulf microbial prey (29), but our study highlights how biofilm exopolysaccharides, such as Psl in *Pseudomonas aeruginosa* can impede this process. This suggests that biofilm-forming microbes may exploit their extracellular matrix not only as a physical barrier but also as a means to immobilize phagocytic predators, limiting their capacity to infiltrate and consume biofilm communities. Such interactions could shape microbial survival strategies in natural environments, promoting the persistence of biofilms under predatory pressure. Additionally, the parallels between amoebae and macrophages in their motility responses underscore the evolutionary conservation of these interactions (30), suggesting that insights from amoeba-biofilm dynamics could inform our understanding of immune evasion mechanisms in pathogenic contexts.

These findings open new avenues for exploring how biofilm matrices influence ecological and host-associated predator-prey dynamics, potentially guiding the development of strategies to disrupt biofilm resilience and improve control of biofilm-associated infections (12). By disrupting the biofilm matrix or reducing its adhesiveness (31), this may restore macrophage motility and improve the effectiveness of immune responses against chronic infections.

In summary, our results emphasize not only the importance of considering the antimicrobial treatment challenges posed by biofilms, but also their capacity to reshape the physical and functional landscape of immune cell movement. This perspective opens new avenues for therapeutic intervention aimed at disrupting the biofilm and restoring effective immune clearance in infected tissues.

## Materials and Methods

### Bacterial Strains, Media, and Growth Conditions

*Pseudomonas aeruginosa* strains and plasmids used in this study are listed as follows:

- PAO1: Prototypic nonmucoid wild-type strain (32).
- PAO1/p_*lac*_*-gfp*: Streptomycin resistance, Tn7-transposon is present in PAO1, with constitutively expressed GFP (33).
- Δ*pelA/p*_*lac*_*-gfp*: Streptomycin resistance, Δ*pelA* carrying the Tn7-transposon (*/*p_*lac*_*-gfp*), with constitutively expressed GFP (22).
- Δ*pslBCD/p*_*lac*_*-gfp*: Streptomycin resistance, Δ*pslBCD* carrying the Tn7-transposon, with constitutively expressed GFP (22).
- Δ*pelA*Δ*pslBCD/p*_*lac*_*-gfp*: Streptomycin resistance, Δ*pelA*Δ*pslBCD* carrying the Tn7-transposon, with constitutively expressed GFP (22).
- Δ*wspF /p*_*lac*_*-gfp*: *wspF* knockout of PAO1 constructed by allelic exchange (22).
- Δ*wspF*Δ*pelA/p*_*lac*_*-gfp*: Pel exopolysaccharide defective *pelA* mutant in Δ*wspF* mutant (22).
- Δ*wspF*Δ*pslBCD/p*_*lac*_*-gfp*: Psl exopolysaccharide defective *pslBCD* mutant in Δ*wspF* mutant (22).
- Δ*wspF*Δ*pelA*Δ*pslBCD/p*_*lac*_*-gfp*: Pel and Psl exopolysaccharides defective *pelA* and *pslBCD* mutant in Δ*wspF* mutant (22).

All strains were cultured in Luria-Bertani medium (Difco, Becton Dickinson and Company, USA) or ABTGC medium (ABT minimal medium supplemented with 2 g L^-1^ glucose and 2 g L^-1^ casamino acids) at 37 °C with shaking at 200 rpm (34). Tn7-GFP plasmid was transformed into bacterial strains to achieve the green fluorescent protein (GFP) expression (22). 50 μg mL^−1^ ampicillin (Sigma-Aldrich, Germany) was used to maintain the plasmid. The bacterial strains were cultured in the growth medium supplemented with 100 μg ml-1 streptomycin for marker selection.

### Fabrication of biofilm cultivation device

As previously described (35), we employed a microfluidic model containing microwell-laden channels for our study. The microfluidic molds were initially silanized with trichloro (1H,1H,2H,2H-perfluorooctyl) silane (Sigma-Aldrich, Germany) overnight in a vacuum desiccator (36, 37). The microfluidic device was fabricated using PDMS (Polydimethylsiloxane), prepared from a Sylgard 184 silicone elastomer kit (Dow Corning, USA). This involved thoroughly mixing the base resin and curing agent in a ratio of 10:1 by weight (38). The layers were assembled and treated with plasma for 5 minutes at high RF level (700 mmtor), then placed in a 70 °C oven for at least 2 hours.

### Cultivation of biofilm in device

*P. aeruginosa* biofilms were cultivated in a microfluidic device using a continuous flow system. Prior to each experiment, the device was disinfected with 75% ethanol, rinsed, and any remaining bubbles were removed. Each channel was then inoculated with 100 μl of bacterial culture (10^6^ cells ml^-1^) using a sterile syringe and needle. Following an initial incubation of 20 mins without flow to allow for bacterial colonization of the microwells, ABTGC medium was continuously supplied at a flow rate of 4 ml hr^-1^ for 12 hrs at 37 °C using a syringe pump. The waste medium was directed into a waste beaker.

### Preparation of Pel and Psl-Coated Microwells

Pel and Psl exopolysaccharides were isolated from their static biofilms of Δ*pslBCD*/p_*lac*_-*yedQ* and Δ*pelA*/p_*lac*_-*yedQ* strains respectively, as previously described (22). Static biofilms were cultivated on LB agar supplemented with appropriate antibiotics at 37 °C. After 16 hrs of cultivation, the biofilms were collected, and the extracellular matrix was separated from bacterial cells via centrifugation and sonication. Extracellular DNA and proteins were removed from the crude matrix extract using ethanol precipitation, CaCl_2_, and proteinase K treatment. The purified Pel and Psl exopolysaccharides were then lyophilized and resuspended in sterile water. The prepared Pel or Psl exopolysaccharides were then evenly coated onto the surfaces of microwells at a concentration of 10 μg ml^-1^ and allowed to dry. The Δ*pelA*Δ*pslBCD*/p_*lac*_-*gfp* strain was then cultivated within these coated microwells to perform further analysis.

### Cell culture and differentiation

Human monocyte U937 cells (CRL-1593.2) were obtained from the American Type Culture Collection (ATCC, Rockefeller, MD) and cultured in a modified Roswell Park Memorial Institute (RPMI 1640) medium (Gibco, #11 875 119, USA) supplemented with 10% FBS at 37 °C in a humidified incubator with 5% CO_2_. Cell passages were performed when cell confluence reached 80%. To differentiate the monocytes into macrophages, U937 monocytes were plated in cell culture dishes at a density of 7 × 10^5^ cells in each dish and treated with 100 ng ml^-1^ phorbol 12-myristate 13-acetate PMA (eBioscience, USA) for 72 hrs. The culture medium containing PMA was renewed every 24 hrs. The PMA-induced M0 macrophages were then stimulated with LPS (eBioscience, USA) at a concentration of 100 ng ml^-1^ for 24 hrs to generate pro-inflammatory M1 macrophages.

### Cultivation of macrophage-biofilm interaction model

The interaction between *P. aeruginosa* biofilms and macrophages was investigated through co-cultivation experiments. Macrophages were prepared by washing with 1X PBS (Gibco), disassociating from dish surface using 0.25% trypsin, phenol red (Gibco), and staining their cell membranes with a Tracking Deep Red Fluorescence dye (Deepred, Invitrogen, USA). A volume of 100 μl of the labelled macrophage suspension (5 × 10^5^ cells ml^-1^) was introduced into each microfluidic channel containing *P. aeruginosa* biofilms. To visualize cell death, propidium iodide (PI, Invitrogen, USA) was added to the culture medium at a final concentration of 20 μM. The biofilms and macrophages were co-incubated at 37 °C and 5% CO_2_ for 2 hrs, with the medium supplied at a flow rate of 4 ml hr^-1^ (similar to diffusion rate) (39) to provide nutrients and remove waste continually. Confocal microscopy was used to capture multiple images (n≥5) from bright-field, deep red fluorescence, and propidium iodide fluorescence channels to observe the interactions between the P. aeruginosa biofilms and macrophages.

### Quantification of cytoskeletal changes and energy utilization

To evaluate cytoskeletal changes in macrophages, F-actin in the macrophages was visualized using 1 μM SiR-Actin (Spirochrome) probe (40) after incubation with SiR-Actin for 40 mins at 37 °C. To assess the energy utilization of the macrophages, 100 nM Tetramethylrhodamine, methyl ester (TMRM, Invitrogen) (41) was used to stain the mitochondria and evaluate ATP usage by the macrophages over 2 hrs. Fluorescent intensities of the SiR-Actin and Tetramethylrhodamine signals were quantified using ImageJ to determine the changes in cytoskeletal organization and energy utilization. The formula used to quantify the cell fluorescence levels was:

Corrected total cell fluorescence = Integrated density - (Area of selected cells × Mean fluorescence of background readings).

### Microscopy imaging and quantitative analysis of macrophages - biofilm interactions

The movement of overall macrophage populations on biofilms were visualized using microscopy video imaging. Microscopic images and videos (every 2 mins) of macrophages on the biofilms were captured by a confocal laser scanning microscope (CLSM) (Leica TCS SP8 MP Multiphoton/Confocal Microscope system) equipped with a 10X objective and an additional 20X zoom to capture brightfield, GFP fluorescence, deepred fluorescence and PI fluorescence in Z-stacks. The images and videos were processed using ImageJ, LAS X, and Imaris software (BitplaneAG, Zurich, Switzerland).

Cell tracking was performed in Imaris software following the manufacturer’s guidelines. The software enabled the tracking of cell movement trajectories and the quantification of macrophage movement metrics, such as average distance and velocities, which were then extracted and exported for further analysis.

### Ex vivo porcine skin infection model

As previously described (42), fresh porcine skin was obtained from a local butcher immediately after slaughter, shaved, and rinsed to remove residual debris. Square skin explants (12 × 12 mm, 5 mm thick) were excised using a sterile scalpel. A central circular wound bed (3 mm diameter, ∼1.5 mm depth) was created in each explant to simulate a partial-thickness skin wound. Explants were sterilized by immersion in 70% ethanol for 30 mins, followed by 10% sodium hypochlorite for 30 mins. They were then rinsed 3 times in sterile 0.9% NaCl (w/v) to remove residual disinfectants. To prevent microbial penetration through the explant base, sterilized explants were placed epidermis-side up on soft tryptic soy agar (TSA; 0.5% agar; NutriSelect, Germany) supplemented with 50 μg/mL gentamicin (Gibco, USA).

For infection, 10 μL of bacterial suspension (1 × 10_ CFU) was applied directly to the wound bed. Explants were incubated at 37 °C in 5% CO_2_ and saturated humidity for 48 hrs to allow biofilm formation. To maintain tissue hydration, explants were transferred daily to fresh soft TSA plates containing 0.5% agar and gentamicin. To examine macrophage–biofilm interactions in the ex vivo model, macrophages were pre-stained with Tracking Deep Red Fluorescence dye (Deepred, Invitrogen, USA). A 200 μL macrophage suspension (5 × 10□ cells/mL) was seeded into a modified 8-well chamber (Ibidi, USA) equipped with two side channels to permit continuous medium flow and waste removal, ensuring stable nutrient conditions.

The 100 μm-thick section of biofilm-infected porcine skin explant was aseptically excised and placed directly onto the macrophage monolayer. Culture medium containing 20 μM propidium iodide was perfused through the chamber at a constant rate of 4 ml/hr via a syringe pump (12). Co-incubation was conducted at 37 °C in a humidified atmosphere with 5% CO_2_ for 2 hrs.

Live imaging of macrophage motility over biofilm-infected skin sections was performed using the CLSM (Leica TCS SP8 MP confocal microscope) equipped with a 10× objective and 20× digital zoom. Z-stack images were captured in brightfield, GFP (biofilm), deep-red (macrophages), and PI (dead cells) channels every 2 min for the duration of the experiment.

### Mathematical Modeling

The mathematical modelling approach involved video processing techniques for cell tracking, followed by data analysis using statistical modelling and simulation methods. Time-lapse microscopy videos were processed using Imaris software, utilizing the “Spots” function to track individual macrophages. A spot diameter of 20 µm (average size of a human macrophage) was selected to encapsulate individual macrophages. Macrophage segmentation from the background was achieved through a background subtraction method, followed by manual thresholding to remove outlier pixels. Filters were then implemented to enhance the contrast between macrophage pixels. Key parameters characterizing macrophage movement, such as velocity and distance in three dimensions, were calculated from the tracked trajectories.

The extracted trajectory data were further analysed using R studio. Exponential smoothing was applied to reduce random fluctuations and normalize the data using the following equation:

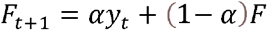

where Ft is the previous smoothed value at time t, Ft+1 is smoothed statistic, α is the smoothing factor (0 < α < 1), and yt is the observed value of series at time t.

A random walk simulation was developed in R Studio. The simulation framework incorporated the biofilm’s matrix density. Individual macrophages were represented as agents within the simulated environment, each with a defined set of movement rules. Polynomial regression models were employed, as follows:

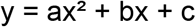

where a, b, and c represent the coefficients used for the modeling.

The distance equations for macrophages on each mutant biofilm were presented as follows:

Macrophages on PAO1 biofilm:

y = -0.000002x^2^ - 0.0016x + 1.949, with an R^2^ value of 0.9954.

Macrophages on Δ*wspF* biofilm:

y = 0.0000009x^2^ + 0.0002x + 0.8905, with an R^2^ value of 0.9592.

Macrophages on Δ*wspF*Δ*pelA* biofilm:

y = 0.0000004x^2^ - 0.0019x + 1.8825, with an R^2^ value of 0.9954.

Macrophages on Δ*wspF*Δ*pslBCD* biofilm:

y = 0.000004x^2^ - 0.0015x + 3.2827, with an R^2^ value of 0.9746.

Macrophages on Δ*wspF*Δ*pelA*Δ*pslBCD* biofilm:

y = -0.00001x^2^ - 0.0012x + 3.6331, with an R^2^ value of 0.9889.

Macrophages on Δ*pelA* biofilm:

y = 0.000006x^2^ - 0.0024x + 2.2201, with an R^2^ value of 0.9956.

Macrophages on Δ*pslBCD* biofilm:

y = 0.00001x^2^ - 0.0062x + 2.6413, with an R^2^ value of 0.9925.

Macrophages on Δ*pelA*Δ*pslBCD* biofilm:

y = 0.000007x^2^ - 0.0037x + 3.0946, with an R^2^ value of 0.9950.

The velocity equations for macrophages on each mutant biofilm were presented as follows:

Macrophages on PAO1 biofilm:

y = -0.000004x^2^ - 0.0015x + 0.7739, with an R^2^ value of 0.9954

Macrophages on Δ*wspF* biofilm:

y = 0.000002x^2^ + 0.0002x + 0.3537, with an R^2^ value of 0.9593.

Macrophages on Δ*wspF*Δ*pelA* biofilm:

y = 0.0000009x^2^ - 0.0018x + 0.7469, with an R^2^ value of 0.9954.

Macrophages on Δ*wspF*Δ*pslBCD* biofilm:

y = 0.00001x^2^ - 0.0014x + 1.294, with an R^2^ value of 0.9746.

Macrophages on Δ*wspF*Δ*pelA*Δ*pslBCD* biofilm:

y = -0.00003x^2^ - 0.0011x + 1.4433, with an R^2^ value of 0.9889.

Macrophages on Δ*pelA* biofilm:

y = 0.00001x^2^ - 0.0023x + 0.881, with an R^2^ value of 0.9956

Macrophages on Δ*pslBCD* biofilm:

y = 0.00003x^2^ - 0.006x + 1.0528, with an R^2^ value of 0.9925

Macrophages on Δ*pelA*Δ*pslBCD* biofilm:

y = 0.00002x^2^ - 0.0036x + 1.2294, with an R^2^ value of 0.9950.

The distance equations for macrophages on each mutant biofilm established on the porcine skin infection model were presented as follows:

Macrophages on PAO1 biofilm established on porcine skin:

y = -0.000007x^2^ - 0.0048x + 2.1388, with an R^2^ value of 0.9662.

Macrophages on Δ*pelA*Δ*pslBCD* biofilm established on porcine skin:

y = 0.000004x^2^ - 0.0012x + 3.6709, with an R^2^ value of 0.9556.

Macrophages on Δ*wspF* biofilm established on porcine skin:

y = 0.000006x^2^ - 0.0059x + 1.8412, with an R^2^ value of 0.9963.

Macrophages on Δ*wspF*Δ*pslBCD* biofilm established on porcine skin:

y = 0.00002x^2^ - 0.0038x + 3.9709, with an R^2^ value of 0.9672.

Macrophages on Δ*wspF*Δ*pelA*Δ*pslBCD* biofilm established on porcine skin:

y = 0.00002x^2^ - 0.0014x + 3.9459, with an R^2^ value of 0.9458.

The velocity equations for macrophages on each mutant biofilm established on the porcine skin infection model were presented as follows:

Macrophages on PAO1 biofilm established on porcine skin:

y = 0.000007x^2^ - 0.0045x + 1.9996, with an R^2^ value of 0.9662.

Macrophages on Δ*pelA*Δ*pslBCD* biofilm established on porcine skin:

y = 0.000004x^2^ - 0.0011x + 3.4319, with an R^2^ value of 0.9555.

Macrophages on Δ*wspF* biofilm established on porcine skin:

y = 0.000005x^2^ - 0.0054x + 1.6748, with an R^2^ value of 0.9963.

Macrophages on Δ*wspF*Δ*pslBCD* biofilm established on porcine skin:

y = 0.00002x^2^ - 0.0036x + 3.7125, with an R^2^ value of 0.9672.

Macrophages on Δ*wspF*Δ*pelA*Δ*pslBCD* biofilm established on porcine skin:

y = 0.00002x^2^ - 0.0014x + 3.6892, with an R^2^ value of 0.9458.

### Statistical analysis

The data were presented as means and standard deviations. Statistical analysis was conducted using one-way ANOVA and Student’s t-tests to evaluate relationships between the independent variables, with p-values reported. Each experiment was performed in triplicate over three independent trials, and the results are reported as the mean ± standard deviation.

## Acknowledgments

This research is supported by The Hong Kong Polytechnic University, Department of Applied Biology and Chemical Technology Departmental General Research Fund (UALB), One-line account (ZVVV), Environmental and Conservation Fund (ECF-84/2021), Health and Medical Research Fund (HMRF-20190302).

## Author Contributions

S.L.C designed methods and experiments. K.M. and Y.M performed laboratory experiments and analyzed the data. K.M., Y.M and S.L.C interpreted the results and wrote the paper. All authors have contributed to, seen, and approved the manuscript.

## Data Availability Statement

The datasets generated during and/or analysed during the current study are available from the corresponding author on reasonable request.

