## Supplemental Figures 1-3 for "Biofilm-derived polymer networks constrain immune cell motility through adhesion-mediated physical interactions"

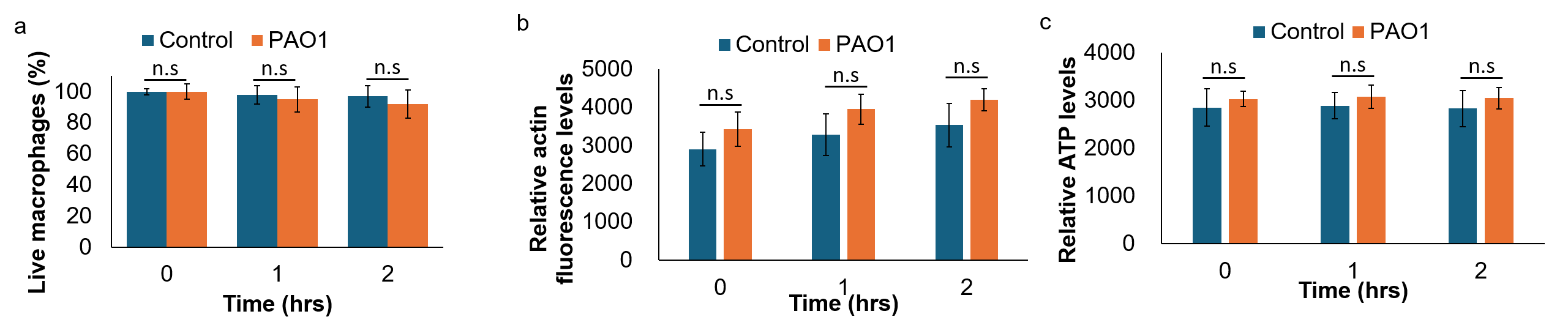
 **Supplementary Figure 1.** Macrophages remain alive and active over time. (a) Minimal death of macrophages, as shown using PI staining. Macrophages are active in actin polarization (b) and ATP usage (c). Means and SD from triplicate experiments in three independent trials are shown. **P < 0.01; ***P < 0.001; n.s: not significant; One-Way ANOVA.

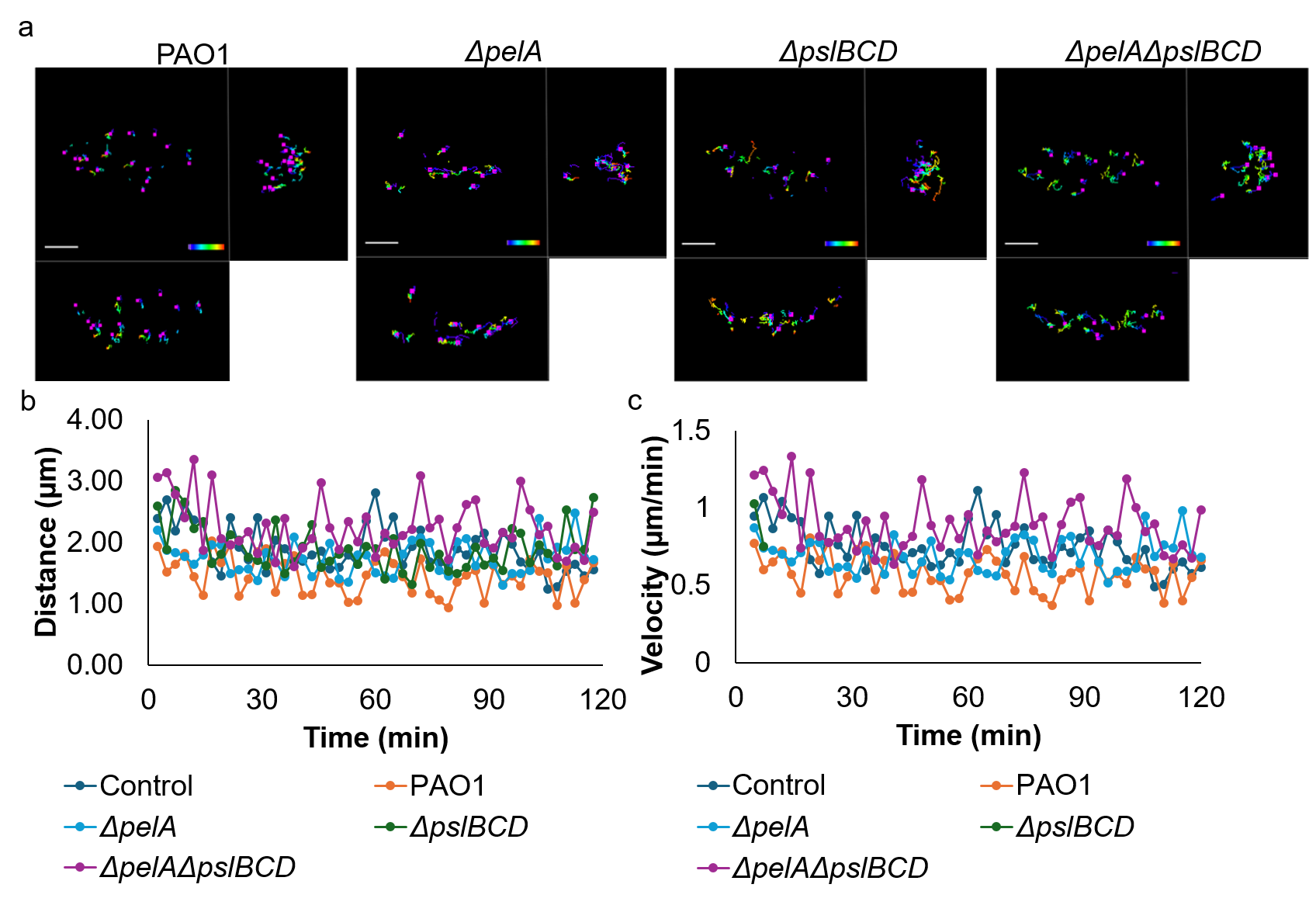

**Supplementary Figure 2.** Exopolysaccharides are important at impeding macrophage motility. (a) Representative tracks, (b) average velocity, and (c) average distance of macrophages on PAO1, Δ*pelA*, Δ*pslBCD*, and Δ*pelA*Δ*pslBCD*.

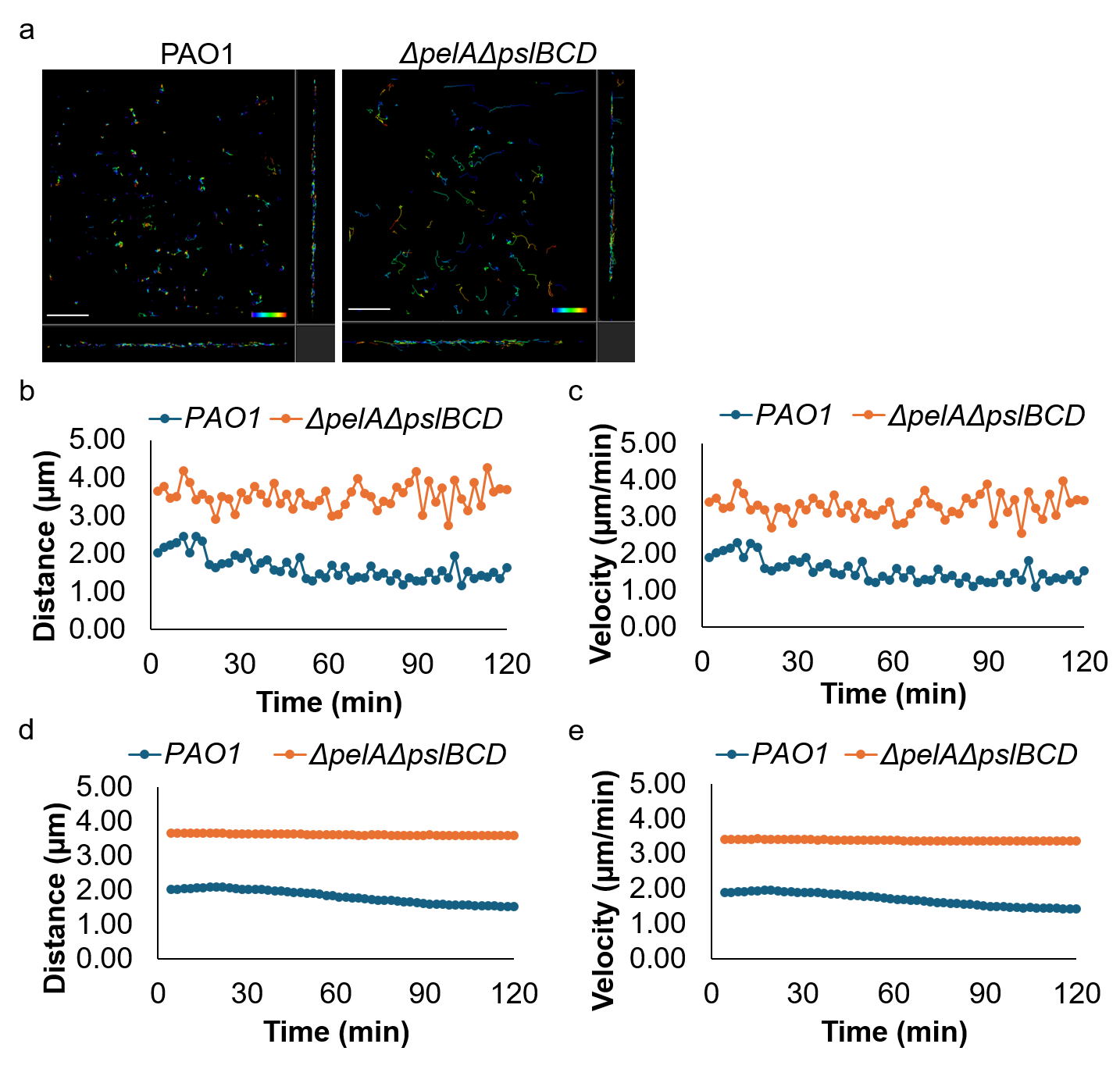

**Supplementary Figure 3.** (a) Representative tracks, (b) average distance, and (c) average velocity of macrophages on PAO1 and Δ*pelA*Δ*pslBCD* pre-established on the porcine skin. Mathematical simulation of (d) relative distance and (e) relative velocity of macrophages motility on various biofilm pre-established on the porcine skin, based on PRW model. Means and SD from triplicate experiments in three independent trials are shown. **P < 0.01; ***P < 0.001; n.s: not significant; One-Way ANOVA.

**Supplementary videos:**

**Video 1.** Motility of macrophages on chip surface.

**Video 2.** Motility of macrophages on PAO1 biofilm.

**Video 3.** Motility of macrophages on Δ*wspF* biofilm.

**Video 4.** Motility of macrophages on Δ*wspF*Δ*pelA* biofilm.

**Video 5.** Motility of macrophages on Δ*wspF*Δ*pslBCD* biofilm.

**Video 6.** Motility of macrophages on Δ*wspF*Δ*pelA*Δ*pslBCD* biofilm.

**Video 7.** Motility of macrophages on Δ*pelA* biofilm.

**Video 8.** Motility of macrophages on Δ*pslBCD* biofilm.

**Video 9.** Motility of macrophages on Δ*pelA*Δ*pslBCD* biofilm.

**Video 10.** Motility of macrophages on Δ*wspF*Δ*pelA*Δ*pslBCD* biofilm with exogenous Pel.

**Video 11.** Motility of macrophages on Δ*wspF*Δ*pelA*Δ*pslBCD* biofilm with exogenous Psl.

**Video 12.** Motility of macrophages on PAO1 biofilm established on the porcine skin infection model.

**Video 13.** Motility of macrophages on Δ*pelA*Δ*pslBCD* biofilm established on the porcine skin infection model.

**Video 14.** Motility of macrophages on Δ*wspF* biofilm established on the porcine skin infection model.

**Video 15.** Motility of macrophages on Δ*wspF*Δ*pslBCD* biofilm established on the porcine skin infection model.

**Video 16.** Motility of macrophages on Δ*wspF*Δ*pelA*Δ*pslBCD* biofilm established on the porcine skin infection model.
